# Causal evidence for the role of dorsolateral prefrontal cortex in the implementation of cognitive control adjustments

**DOI:** 10.64898/2026.09.26.754661

**Authors:** Jacob Sellers, Han Zhang, John Jonides, Taraz Lee

## Abstract

Cognitive control processes enable us to use recent contextual information to flexibly adapt to current task demands. This ability to adapt control processes is exhibited in response conflict tasks. In these tasks, participants must respond to a relevant piece of information while overcoming the tendency to respond to task-irrelevant information that cues an automatic or habitual response. Control demands can be updated in these tasks via two separate routes. Both errors and conflict between task-irrelevant and task-relevant information affect performance on subsequent trials. A body of work has implicated dorsolateral prefrontal cortex (DLPFC) in implementing these control adjustments. However, this work has primarily relied on correlational techniques or focused on a single index of control adaptation. Here, we combine inhibitory transcranial magnetic stimulation (TMS) over DLPFC with computational modeling of behavior to test the causal role of DLPFC in control adaptation. We show that inhibitory TMS to DLPFC reduces the effect of previous trial conflict, and committing an error on the previous trial further reduces adaptive effects. This result provides causal evidence for DLPFC playing a general role in implementing trial-to-trial control adjustments.

## Introduction

Cognitive control allows us to engage in goal-directed action when our goals conflict with our habits (Badre, 2025). For example, resisting the urge to check a phone notification during an important meeting requires cognitive control. Response conflict tasks are often used to investigate this conflict resolution process. In these tasks, participants are instructed to respond to task-relevant information while ignoring often habitual but task-irrelevant information. Not only is performance on any single trial in these tasks dependent on processing relevant information and ignoring irrelevant information, but it also depends on what transpired on a previous trial. For example, experiencing response conflict or committing an error can affect behavior on subsequent trials (Gratton et al., 1992; Rabbitt, 1966). These trial history effects are often used to investigate how we flexibly adapt control processes depending on recent context. This investigation has suggested that the lateral prefrontal cortex (LPFC) plays a key role in implementing flexible adaptations of cognitive control (Carter & Van Veen, 2007; Miller & Cohen, 2001).

Evidence for the role of LPFC in adaptive control has primarily come from neuroimaging studies in humans. These studies have identified distinct brain regions that support various components of cognitive control, including the posterior medial frontal cortex (PMFC) and the LPFC (Assem et al., 2020; Nee et al., 2007, 2011). Neurocomputational theories of control propose that PMFC evaluates and signals a need for control while LPFC responds to this signaling and implements the actual adjustments in control (Botvinick et al., 2001, 2004; Carter & Van Veen, 2007; Kerns et al., 2004; Ridderinkhof et al., 2004; Shenhav et al., 2013).

Two classic indices of control adjustment are the congruency sequence effect (CSE) and post-error effects. In response conflict tasks, the CSE is the finding that the congruency effect, namely the difference in response time between incongruent and congruent trials, is smaller after an incongruent trial than after a congruent trial (Egner, 2007; Gratton et al., 1992). Post-error effects include slowing of response times after making an error (Laming, 1979; Rabbitt, 1966), and reductions in the congruency effect after making an error (Ridderinkhof, 2002).

Prior work has shown increased LPFC activity following incongruent trials and errors (Danielmeier et al., 2011; Egner & Hirsch, 2005; Kerns et al., 2004). However, this work relies on inherently correlational methods, such as functional neuroimaging, leaving the causal contribution of LPFC unclear. Noninvasive brain stimulation techniques, such as transcranial magnetic stimulation (TMS), provide a way to causally test the role of LPFC in cognitive control (Friehs et al., 2020; Lee & D’Esposito, 2012; Muhle-Karbe et al., 2018; Zanto et al., 2011). To date, few studies have used TMS to test the causal role of DLPFC in adaptive control (Friehs et al., 2020; Muhle-Karbe et al., 2018). These studies have primarily examined congruency sequence effects, making it difficult to determine the causal role of LPFC in control adjustments driven by different sources of information. Additionally, almost all studies examining the neural underpinnings of adjustments in cognitive control rely on free response time paradigms. Free response time paradigms do not reveal the temporal dynamics of cognitive control because they measure just the outcome of that control, making it difficult to determine if an adjustment in cognitive control affects task-irrelevant processing, task-relevant processing, or a combination of both. To address this issue, one can use the forced-response method to trace the temporal dynamics of cognitive control in response conflict tasks (Lee et al., 2024). Paired with a computational model of response preparation, data from this method can be used to identify the target of control processes (Sellers et al., 2026).

We applied inhibitory transcranial magnetic stimulation to the right dorsolateral prefrontal cortex (DLPFC) to assess the causal role of DLPFC in adaptations of cognitive control while participants performed a forced-response conflict task. We find evidence that the magnitude of the CSE decreases after DLPFC stimulation relative to stimulation at a control site. This dampening effect of the CSE was especially pronounced when the previous trial was an incorrect trial compared to when it was correct. Computational modeling shows that inhibitory stimulation of DLPFC reduces adjustments in task-irrelevant, habitual processing and task-relevant, goal-directed processing. To our knowledge, these results represent the first causal evidence that DLPFC supports flexible adjustments in cognitive control by modulating the processing of task-irrelevant and task-relevant information to resolve future response conflicts.

## Methods

### Sample

A sample of 21 individuals was recruited to participate in the study. Although we intended to include only right-handed individuals in the study given the laterality of stimulation, one left-handed individual was erroneously tested as well. This individual was excluded from the analysis, leaving a final sample size of 20 (n=12 female, mean age=24.5 years). Participants were recruited from the University of Michigan and the local community. All participants gave informed consent to participate and underwent TMS and fMRI safety screening. Approval was obtained from the Institutional Review Board of the University of Michigan.

### Design

Participants came to the lab for a total of 3-4 in-person sessions and completed one session at home. The first session was used for determining individualized TMS parameters (see TMS protocol). Sixteen participants additionally underwent a structural MRI and a resting-state fMRI on the same day as the individualized TMS session. Another four participants completed the scans on separate day than the initial TMS session due to scheduling conflicts. In the second session, participants were given access to the task online for practice in preparation for the TMS sessions. They completed this practice session from their personal computers via Pavlovia, a site to host psychology experiments online (Peirce et al., 2019). The average time between completion of the online practice and first TMS session was 2 days and 16 hours (*range:* 2 hours–27 days and 23 hours). The online practice session was the same task that was completed in-person (see *Task* below). Following completion of online practice, participants returned for two sessions in which inhibitory TMS was applied to either DLPFC or a control site (see *TMS Protocol* below). An active control site was used to account for any non-specific effects of stimulation on behavior (e.g., auditory sounds, cutaneous stimulation, total current induced in the brain, etc.). In the two TMS sessions, participants completed a forced-response version of the Simon task (see *Task* below). Each task began with 40 practice trials of the Simon task with a free-response procedure, 60 trials of timing practice, and 10 blocks of 60 forced-response task trials (see below for forced-response protocol). TMS was administered after the two practice blocks, and participants began the task trials immediately following the delivery of TMS. The task trials took ∼45 minutes to complete which is well within the ∼60-minute window that the effects of our stimulation protocol is thought to last (Huang et al., 2005). This allowed for an assessment of performance on the Simon task during the period of disruption. Both in-person TMS sessions involved the identical protocol aside from the site of stimulation. The stimulation site order was counterbalanced across participants. The average time between the two in-person TMS sessions was 5 days and 7 hours (*range:* 22 hours–33 days and 20 hours).

### Task

Participants performed a version of the Simon task, which is a task classically used to study the effect of stimulus-response interference (Simon et al., 1976). In this variant of the Simon task, the goal is to respond to the direction of an arrow (Figure 1A). The arrow can point either to the right or to the left. If the arrow was pointing to the left, participants were instructed to press the ‘w’ key on a standard keyboard with their left index finger. If the arrow was pointing to the right, participants were instructed to press the ‘p’ key with their right index finger. The location of the arrow was presented either to the left or to the right of a central fixation stimulus (0.33 height units to the left or right), leading to conflict if the location of the arrow was opposite to its direction. Trials on which there is a mismatch between location and direction of the arrow are referred to as incongruent trials (e.g., arrow pointing to the right on the left side of the screen), and trials on which the location of the arrow is aligned with the direction are referred to as congruent trials (e.g., arrow pointing to the right on the right side of the screen). Each block contained an equal proportion of incongruent and congruent trials (60 per block).

**Figure 1.**
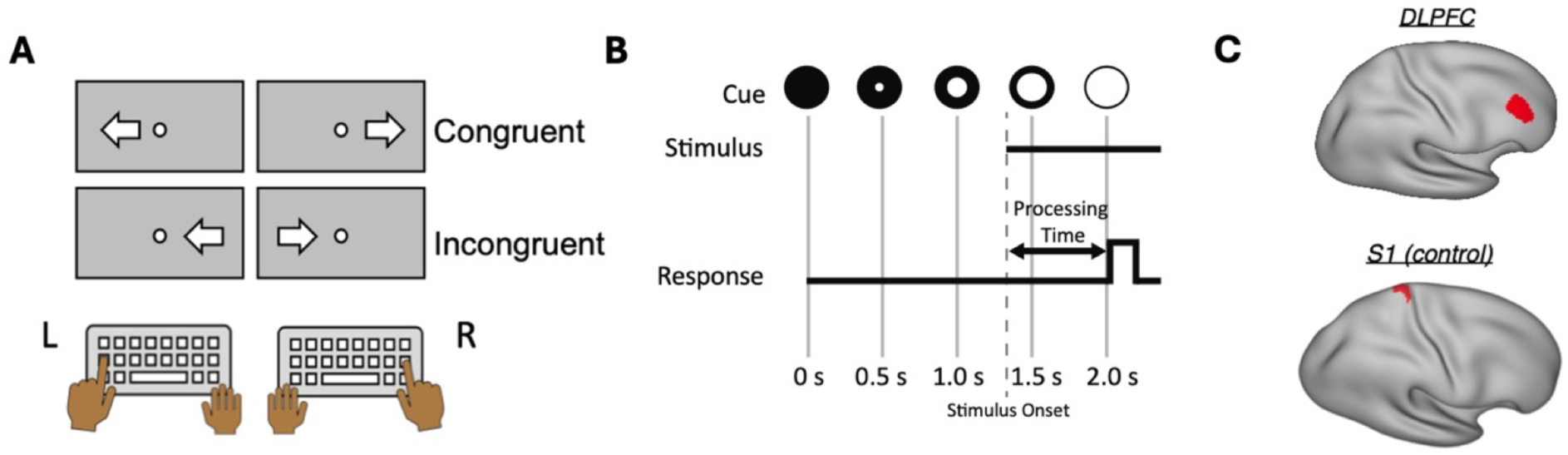
Behavioral paradigm and stimulation sites. *A)* The arrow Simon task. Participants are instructed to respond to the direction of the arrow while ignoring its location. The center circle served as the timing cue in the forced-response method. *B)* The forced-response method. Participants are instructed to respond when the filling circle (cue) turns completely white. Stimulus onset time is varied from trial to trial, allow for processing time to be varied. *C)* DLPFC and S1 (control) stimulation regions.

After training on the Simon task and after practicing the timing of their responses as implemented in the forced-response method, participants performed a forced-response version of the Simon task (Lee et al., 2024). With the forced-response method, response time is fixed, and stimulus onset is varied, giving control over how much time participants have to process stimulus information. A schematic illustration of the method is shown in Figure 1B. Participants were instructed to respond when a centrally placed circle completely turned white. The circle gradually filled over time (25% every 500 ms) and was completely filled 2000 ms from the start of every trial. The time of stimulus onset was randomly drawn from a uniform distribution ranging from 1000-2000 ms, meaning that the time between stimulus onset and the cue to respond varied between 0-1000 ms. Naturally, participants most often did not respond at the exact time they were cued, so we define “processing time” as the difference between the actual response time and stimulus onset on each trial. For example, if the stimulus appeared at 1500 ms and the participant responded at 2050 ms, processing time was defined as 550 ms for that trial. Participants were given feedback on their timing performance after each trial. If their response came more than 100 ms prior to the timing cue, they received a message stating, “Too fast!”. If their response came more than 100 ms after the timing cue, they received a message stating, ‘Too slow!’. If they responded within the ±100 ms window relative to the timing cue, they received a message stating, “Perfect timing!”. The critical dependent variable in this task was accuracy since processing time was the independent variable that we sought to systematically vary. This manipulation allowed us to investigate how accuracy changes over time (i.e., the speed-accuracy tradeoff function) using our response-preparation model.

### fMRI Preprocessing

MR data were acquired with a 3T GE scanner (MR 750) with a 32-channel head coil. Functional data were obtained using a 1-shot multi-band T2∗-weighted echo-planar imaging (EPI) sequence sensitive to blood oxygenation level-dependent (BOLD) contrast (TR = 650 ms, TE = 34.8 ms, flip angle = 54 °, 2.3 mm isotropic voxels, multiband factor = 8). Whole-brain T1-weighted scans were acquired for anatomical localization. Resting-state fMRI data were first preprocessed using fMRIPrep (Version 20.2.5). First, a reference volume and its skull-stripped version were generated using the custom methodology of fMRIPrep. A B0-nonuniformity map (or fieldmap) was estimated based on two (or more) echo-planar imaging (EPI) references with opposing phase-encoding directions, with 3dQwarp (Cox & Hyde, 1997; AFNI 20160207). Based on the estimated susceptibility distortion, a corrected EPI reference was calculated for a more accurate co-registration with the anatomical reference. The BOLD reference was then co-registered to the T1w reference using bbregister (FreeSurfer), which implements boundary-based registration (Greve & Fischl, 2009). Co-registration was configured with six degrees of freedom. Head-motion parameters with respect to the BOLD reference (transformation matrices, and six corresponding rotation and translation parameters) were estimated before any spatiotemporal filtering using mcflirt (FSL 5.0.9; Jenkinson et al., 2002). The BOLD time-series were resampled onto their original, native space by applying a single, composite transform to correct for head-motion and susceptibility distortions. Several confounding time-series were calculated based on the preprocessed BOLD: framewise displacement (FD) and three region-wise global signals. FD was computed using the Power formulation (absolute sum of relative motions, Power et al., 2014). The three global signals were extracted within the cerebrospinal fluid (CSF), the white matter (WM), and the whole-brain masks. The confound time series derived from head motion estimates and global signals were expanded with the inclusion of temporal derivatives and quadratic terms for each (Satterthwaite et al., 2013). Frames that exceeded a threshold of 0.2 mm FD were considered motion outliers. All re-samplings can be performed with a single interpolation step by composing all the pertinent transformations (i.e. head-motion transform matrices, susceptibility distortion correction when available, and co-registrations to anatomical and output spaces). Gridded (volumetric) re-samplings were performed using antsApplyTransforms (ANTs), configured with Lanczos interpolation to minimize the smoothing effects of other kernels (Lanczos, 1964). Following fMRIPrep, we performed band-pass filtering and confound regression using the nilearn python package. A Butterworth temporal filter extracted frequencies between 0.009 and 0.08 Hz. The band-pass filter was also applied to all confound regressors to avoid reintroducing filtered frequency bands (Lindquist et al., 2019). The confound regression included the six head motion estimates and three region-wise global signals (CSF, WM, whole-brain), their temporal derivatives, quadratic terms, and the quadratic expansions of their derivatives (36 regressors; Satterthwaite et al., 2013). We additionally included spike regressors associated with motion outliers. Following confound regression, the resulting residual timeseries were z-scored. We then performed spatial smoothing with a cortical ribbon constrained 4 mm FWHM Gaussian kernel.

### TMS Protocol

Repetitive TMS (rTMS) was delivered through a MagPro X100 magnetic stimulator and a 70 mm figure-8 coil (MC-B70, MagVenture Inc.). In the initial screening session, motor evoked potentials (MEP) elicited using biphasic posterior-anterior stimulation and coil oriented 45 degrees to coronal plane were recorded from left first dorsal interosseous (FDI) using surface electromyography (2-channel built in EMG device, Rogue Research, Montreal CA). Active motor threshold (AMT) was determined as the percentage of stimulator output that elicits an MEP of ≥ 200 μV peak-to-peak on five out of ten trials while the FDI muscle is contracted at 20% of maximum. At the end of this screening session, participants were given several pulses at 80% AMT over regions of the scalp roughly corresponding to DLPFC and the control site to ensure that the stimulation intensity would be tolerable in future sessions. No participants reported discomfort, nor were any participants screened out in this process. In the experimental sessions following training, we delivered continuous theta-burst stimulation (cTBS) over a single region (DLPFC or the control site) with standard parameters and within consensus safety recommendations (Oberman et al., 2011; Rossi et al., 2021): 3 pulses of stimulation at 50 Hz at 80% AMT, repeated every 200 ms, for a total of 600 pulses in 40 seconds. The cTBS protocol has been found to have an inhibitory effect on the site of stimulation for approximately 60 minutes following stimulation (Huang et al., 2005).

### Stimulation Sites

We applied cTBS to either the right DLPFC or a region of somatosensory cortex (S1) prior to performance on a Simon task (Figure 1C). Individual targets were chosen using resting-state functional connectivity to a region implicated in the conflict-resolution process in the right lateral parietal cortex. Anatomical masks were defined using the Human Connectome Project multimodal parcellation (Glasser et al., 2016). We extracted the average time-course from a right lateral parietal (IP2) region of interest and then correlated this time-course with the rest of the brain. An anatomical mask was defined at p46-v, and the maximum correlation to IP2 within this mask was chosen as the DLPFC target. This DLPFC parcel has been implicated in the conflict-resolution process (Freund et al., 2021) and domain-general cognitive control (Assem et al., 2020). An anatomical mask was defined as a medial region in the postcentral gyrus, and the minimum absolute correlation to IP2 was chosen as the control target. This control region approximately corresponded with the lower limb representation of primary somatosensory cortex (S1).

### Behavioral Analysis

Due to natural variations in participants’ response timing, the processing time used in analysis was the actual time between stimulus onset and response time. We excluded trials from analysis that had a processing time of less than 0ms, as these are trials in which the participant responded before the stimulus appeared on the screen (1.06% of trials). We also excluded trials in which processing time exceeded 1000 ms to exclude trials in which participants excessively delayed a response past intended processing times (1.98% of trials).

We visualize the response accuracy as a function of PT using a sliding window procedure in which accuracy was iteratively computed in a 120 ms window, sliding from 0 to 1000 ms with a step size of 1 ms. At each time step, the mean accuracy was computed for each participant in each condition. We averaged participant time courses to obtain group-level time courses. Error bars on these group-level time courses denote standard error. We display the 0 to 700 ms range, as participants perform at near ceiling accuracy after 700 ms. To visualize trial-to-trial effects as a function of stimulation, trajectories were computed for congruent and incongruent trials split by previous trial congruency (congruent and incongruent), previous trial accuracy (correct and error), and stimulation condition (DLPFC and control).

### Response Preparation Model

We conducted a model-based analysis of the forced-response data in which we treat response accuracy arising from competitive preparation of habitual and goal-directed responses (see Figure 2A for a schematic of the model). According to this model, habitual responses to the location of the arrow and goal-directed responses to the direction of the arrow are prepared independently and in parallel, with normally distributed latencies (response-preparation distributions in Figure 2A). Response accuracy is assumed to be a Bernoulli variable with probability *θ*. This probability is assumed to be a function of time *t* and denotes the probability of executing the correct response at *t*. It is defined as the product of a matrix specifying the preparatory state at *t* (*ϕ*) and a matrix of the conditional mappings of preparatory states to responses (*ψ*). Note the subscript *c* and *i* refer to the predictions for congruent and incongruent trials, respectively:

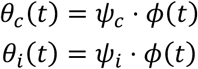

**Figure 2.**
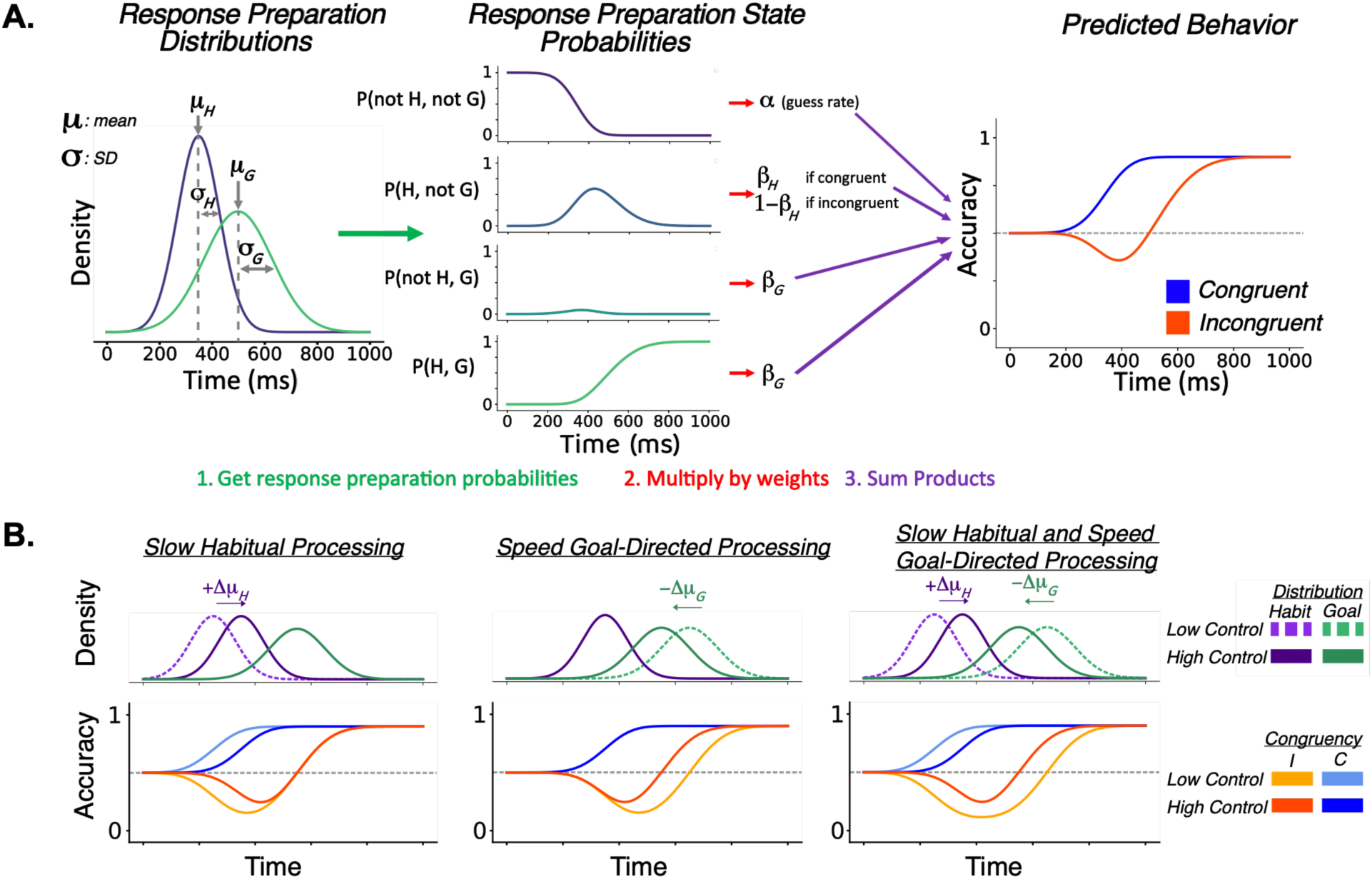
The response preparation model. *A)* Schematic showing how estimated response preparation distributions are translated to behavior. Habitual and goal-directed response preparation latency is modeled as a normal distribution. The probability of being in each response preparation state over time is computed as the joint probability of each response being prepared or not prepared. These response preparation states are multiplied by weights that capture the probability of correctly responding given that one is in some response preparation state. These weights can be thought of as control parameters that govern the likelihood of emitting a response if it is prepared. The weighted probabilities are summed to obtain predicted behavior for both congruent and incongruent trial types. *B)* Alternative hypotheses for changes in cognitive control. Increases in cognitive control allocation (e.g., due to committing an error, experiencing conflict, or TMS) could result in a slowing of habitual response preparation, speeding of goal-directed response preparation, or a combination of both possibilities.

The matrix *ϕ*(*t*) gives the probabilities of all combinations of habitual and goal-directed responses being prepared and unprepared at *t* (response preparation state probabilities in Figure 2A). These probabilities are the probability that nothing is prepared, the habitual response is prepared but the goal-directed response is not prepared, the habitual response is not prepared but the goal-directed response is prepared, and both responses are prepared. These probabilities depend on *t* and the preparation parameters (*μ* and σ):

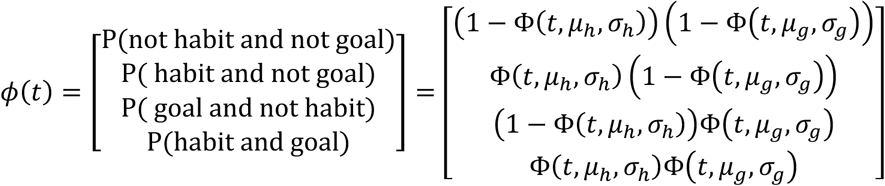

Where Φ denotes the cumulative distribution function for the normal distribution. Then, for example, Φ(*t*, *μ*_ℎ_, *σ*_ℎ_) is the probability of the habitual response being prepared at *t*, and 1 − Φ(*t*, *μ*_ℎ_, *σ*_ℎ_) is the probability of the habitual response not being prepared at *t*. As we assume each response is prepared independently, *ϕ*(*t*) contains the joint probabilities of all combinations of habitual and goal-directed responses being prepared and unprepared.

The matrix *ψ* specifies the probabilities of expressing a habit or goal depending on the preparatory state (weights in Fig. 2a):

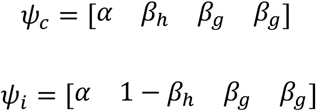

*α* is the probability of expressing the correct response (the goal) if neither the goal nor the habit is prepared. This probability can be thought to capture ‘chance’ performance in a task. *β*_ℎ_ is the probability of expressing the habitual response given that only the habitual response is prepared. Since the habitual response is incorrect on incongruent trials and we are modeling accuracy, we use 1 − *β*_ℎ_ for incongruent trials. *β*_g_ governs the probability of correctly expressing the goal-directed response any time the goal-directed response is prepared. This includes time periods when the goal is prepared but the habit is not prepared, and time periods when both the habit and the goal are prepared. *β*_g_ captures ceiling-level accuracy in forced-response tasks.

In summary, the model yields 7 group-level parameters. These are the means and standard deviations of the habit and goal preparation distributions (*μ*_ℎ_, *μ*_g_, *σ*_ℎ_, *σ*_g_), the probability of emitting the correct response if nothing is prepared *α*, the probability of emitting the habitual response if only the habit is prepared *β*_ℎ_, and the probability of emitting the goal-directed response if the goal is prepared *β*_g_.

The model described above was programmed with the Python library PyMC (Abril-Pla et al., 2023). *α* was fixed at 0.5 during model fitting to increase identifiability. All free parameters were modeled hierarchically using non-centered parameterizations. To control for order effects, all model parameters included a regression effect of session number (session 1 vs session 2). For participant *j* on trial *i*, parameters without additional regressors (*σ*_ℎ_, *σ*_g_, *β*_ℎ_, *β*_g_) were modeled as:

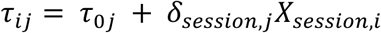

where *τ*_0j_ denotes the participant intercept, *δ_session,j_* denotes the participant-specific session effect, and *X_session,i_* denotes the session predictor.

To test how stimulation and trial history affect the preparation of habitual and goal-directed response preparation, the mean preparation latency parameters *μ*_ℎ_ and *μ*_g_included additional regressors. These regressors captured the effects of stimulation (DLPFC vs control), previous trial congruency (previous incongruent vs previous congruent), previous trial accuracy (previous error vs previous correct), and all lower-and higher-order interactions among these predictors. For *p* ∈ {*μ*_ℎ_, *μ*_g_}:

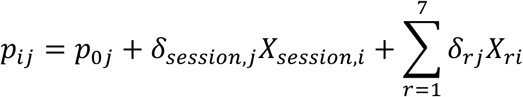

All intercepts and regression coefficients were estimated hierarchically. Participant-level intercepts and coefficients were modeled using non-centered parameterizations:

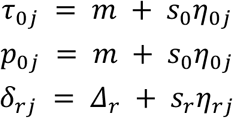

where *m* denotes the group-level intercept (*μ*_ℎ_, *μ*_g_, *σ*_ℎ_, *σ*_g_, *β*_ℎ_, *β*_g_), *Δ_r_* denotes the group-level mean effect, *s_r_* denotes the group-level standard deviation across participants, and *η_rj_* denotes the standardized participant-level deviation. All parameters were sampled in unconstrained space and transformed to the [0,1] interval via the inverse logit function. Priors on group-level intercepts were:

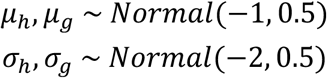

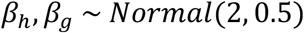

All group-level regression effects were assigned weakly informative priors centered on 0:

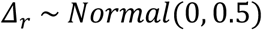

Group-level standard deviations were assigned HalfNormal priors:

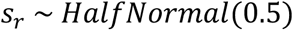

Standardized participant-level deviations were assigned standard normal priors:

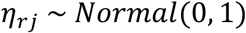

These priors are weakly informative and have been used in past applications of the response preparation model (Osborne et al., 2026; Sellers et al., 2025, 2026). The estimated effects of stimulation, previous trial congruency, and previous trial accuracy were assigned prior distributions centered on zero to reflect ambiguity about these effects. Posterior distributions were estimated using Markov chain Monte Carlo with four chains. Each chain ran for 3000 warm-up iterations and 3000 sample iterations. Convergence was assessed using *R*^L^ and visual inspection of trace plots. All parameter had *Ȓ* < 1.01.

Increases in cognitive control allocation can result in a slowing of habitual processing, speeding of goal-directed processing, or a combination of both possibilities. These different control adjustments would manifest in distinct behavioral patterns, and the response preparation model can distinguish between them (Figure 2B). We obtained an estimate of habitual and goal-directed response preparation speed using estimated group-level intercepts and effects for each of the 8 conditions combinations (i.e., each combination of stimulation, previous trial congruency, and previous trial accuracy). We report the main effect of stimulation, which captures how DLPFC stimulation affects response preparation on the current trial collapsed across trial history effects. We report the effect of previous trial congruency (i.e., the congruency sequence effect), previous trial accuracy (i.e., post-error effect), and the interaction between previous trial congruency and previous trial accuracy for control stimulation as a baseline. We additionally report how DLPFC stimulation alters the CSE, the post-error effect, and the interaction between the CSE and the post-error effect. We report these effects for habitual response preparation latency *μ*_ℎ_ and goal-directed response preparation latency *μ*_g_. We also report these effects for the difference in goal-directed and habitual response preparation latency *μ*_g_ − *μ*_ℎ_. A smaller difference can be interpreted as stronger cognitive control overall, as a smaller difference indicates less time when a habitual response can be expressed in the absence of a goal-directed response. A smaller difference results in a reduced congruency effect (see right panel of Figure 2B).

For each estimated effect, we report the posterior mean, 95% credible interval, and probability of direction (*pd*). The *pd* is the proportion of the obtained posterior distribution that is the same direction as the median of the posterior. The *pd* can range from 0.5 to 1. A *pd* of 0.5 indicates no evidence for the existence of an effect (i.e., a posterior completely centered on 0). A pd close to 1 indicates very strong evidence for the existence of an effect (i.e., a posterior that is almost all negative or all positive). We interpret pd ≥ 0.95 strong evidence for an effect, 0.85 ≤ *pd* < 0.95 as some evidence for an effect, and pd < 0.85 as weak evidence for an effect.

## Results

To preview our results, following control stimulation, we observe a CSE that is larger after error trials than after correct trials. Stimulation of DLPFC reduces the overall CSE, and the CSE is further reduced or even abolished after an error. Computational modeling reveals that these adjustments in control affect both habitual and goal-directed response preparation, and DLPFC stimulation reduces these control adjustments. Behavioral data from the forced-response method reveal the time course of cognitive control (Figure 3). In each condition, accuracy rates are near chance-level performance (50%) until roughly 200 ms before accuracy diverges for congruent and incongruent trial types. Notably, the CSE is visually different as a function of stimulation condition and previous trial accuracy. After a correct trial, the CSE appears weaker following DLPFC stimulation from roughly 200-300 ms (see Figure 3A and 3B). Following an error, the CSE appears even weaker or is abolished following DLPFC stimulation (see Figure 3C and 3D).

**Figure 3.**
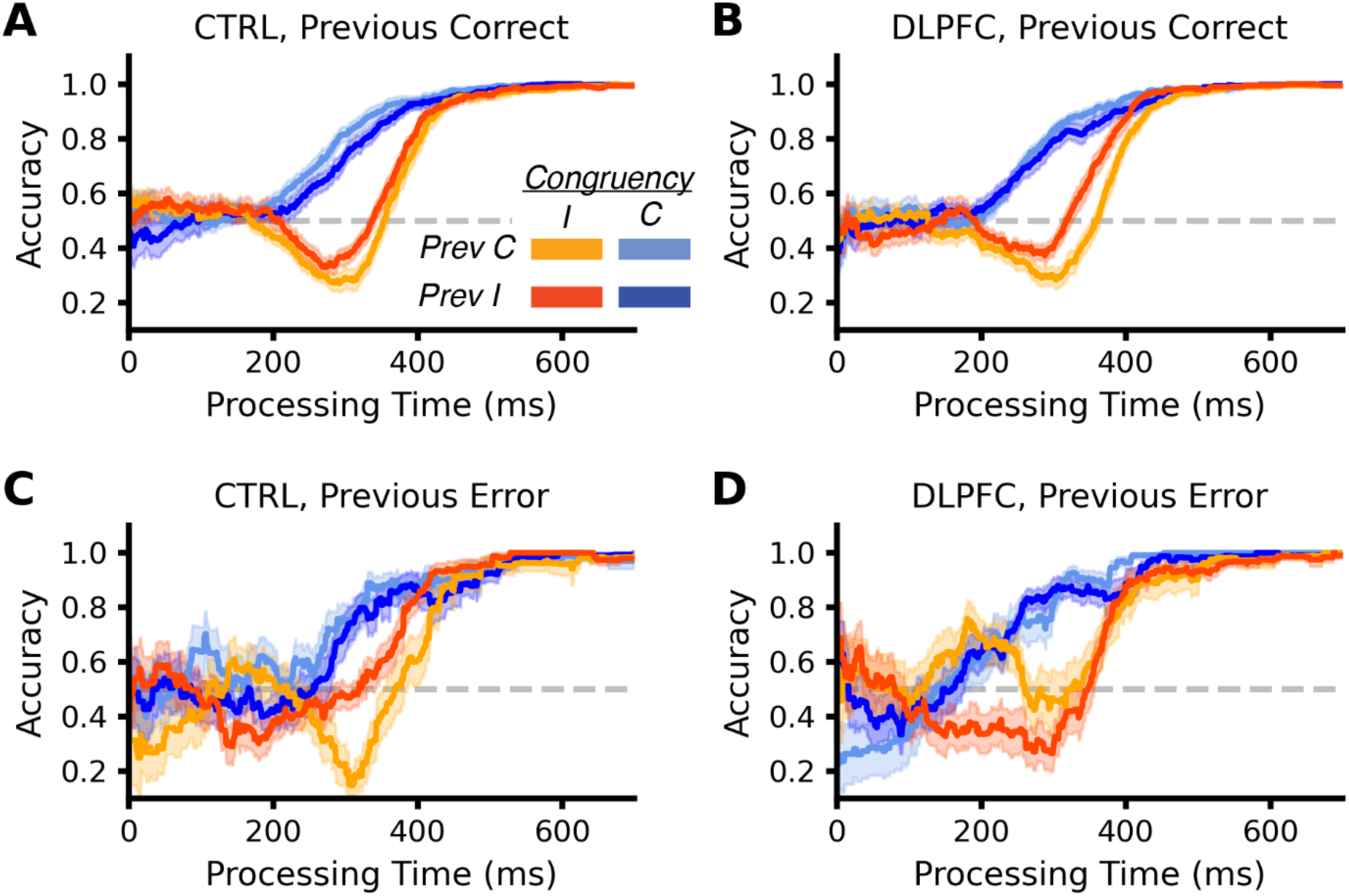
Behavioral data (accuracy as a function of processing time) split by previous trial congruency, previous trial accuracy, and stimulation condition. In all conditions, participants are guessing until roughly 200 ms before a time-dependent congruency effect emerges. Accuracy reaches asymptote by 600 ms in all conditions. *A)* The CSE for control stimulation following a correct response. *B)* The CSE for DLPFC stimulation following a correct response. *C)* The CSE for control stimulation following an error. *D)* The CSE for DLPFC stimulation following an error.

To quantify these effects of stimulation on adjustments in the component processes of cognitive control, we fit a computational model of response preparation to these data (Figure 4). We begin with the effects of stimulation and trial history on habitual response preparation (i.e. processing of task-irrelevant location information; Figure 5, purple). Following control (S1) stimulation, we found strong evidence that habitual responses were prepared more slowly after an incongruent trial (*b* = 24.8 ms, *95CI* = [-1.66, 50.8], *pd* = 0.968). We found weak evidence for a difference in habitual response preparation after error trials (*b* = 5.86 ms, *95CI* = [-19.6, 33.1], *pd* = 0.672), and weak evidence for an interaction between the CSE and post-error effect (*b* = -2.93 ms, *95CI* = [-53.8, 48.3], *pd* = 0.546). We found weak evidence for a main effect of DLPFC stimulation on habitual response preparation (*b* = -8.23 ms, *95CI* = [-28.3, 11.3], *pd* = 0.796). However, we found strong evidence for an interaction between stimulation site and the CSE (*b* = -34.6 ms, *95CI* = [-70.5, 1.12], *pd* = 0.972), suggesting that DLPFC stimulation reduced the slowing of habitual processing following an incongruent trial. We found weak evidence for an interaction between the stimulation and the post-error effect (*b* = -14.9 ms, *95CI* = [-51.2, 21.2], *pd* = 0.790). We found moderate evidence for a three-way interaction between stimulation site, the CSE, and the post-error effect (*b* = -43.1 ms, *95CI* = [-115, 28.0], *pd* = 0.889), suggesting that an error on the previous trial further reduced or even reversed any slowing of habitual response preparation after an incongruent trial for DLPFC stimulation.

**Figure 4.**
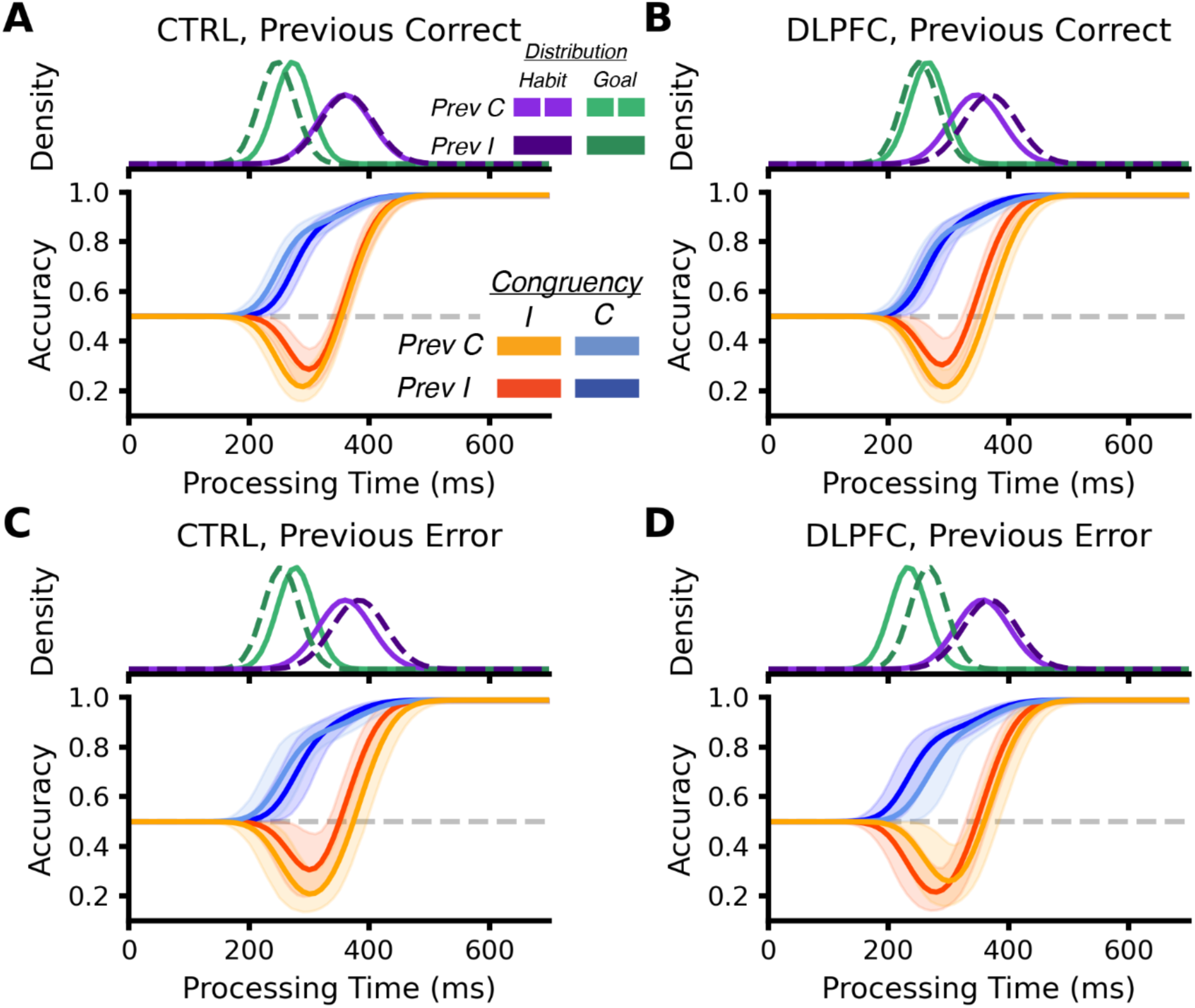
Posterior predictions for behavior and response preparation distributions obtained from fitting the response preparation model. The model predictions demonstrate that the model fit captures key aspects of the behavioral data, namely the time-dependent congruency effect. The model can additionally capture differences in accuracy as a function of processing time between previous trial congruency, previous trial accuracy, and stimulation conditions. Importantly, this approach allows one to estimate the effect of previous trial congruency, previous trial accuracy, and stimulation on habitual and goaldirected response preparation latency (sub panel above each behavioral prediction). *A)* The CSE for control stimulation following a correct response. *B)* The CSE for DLPFC stimulation following a correct response. *C)* The CSE for control stimulation following an error. *D)* The CSE for DLPFC stimulation following an error.

**Figure 5.**
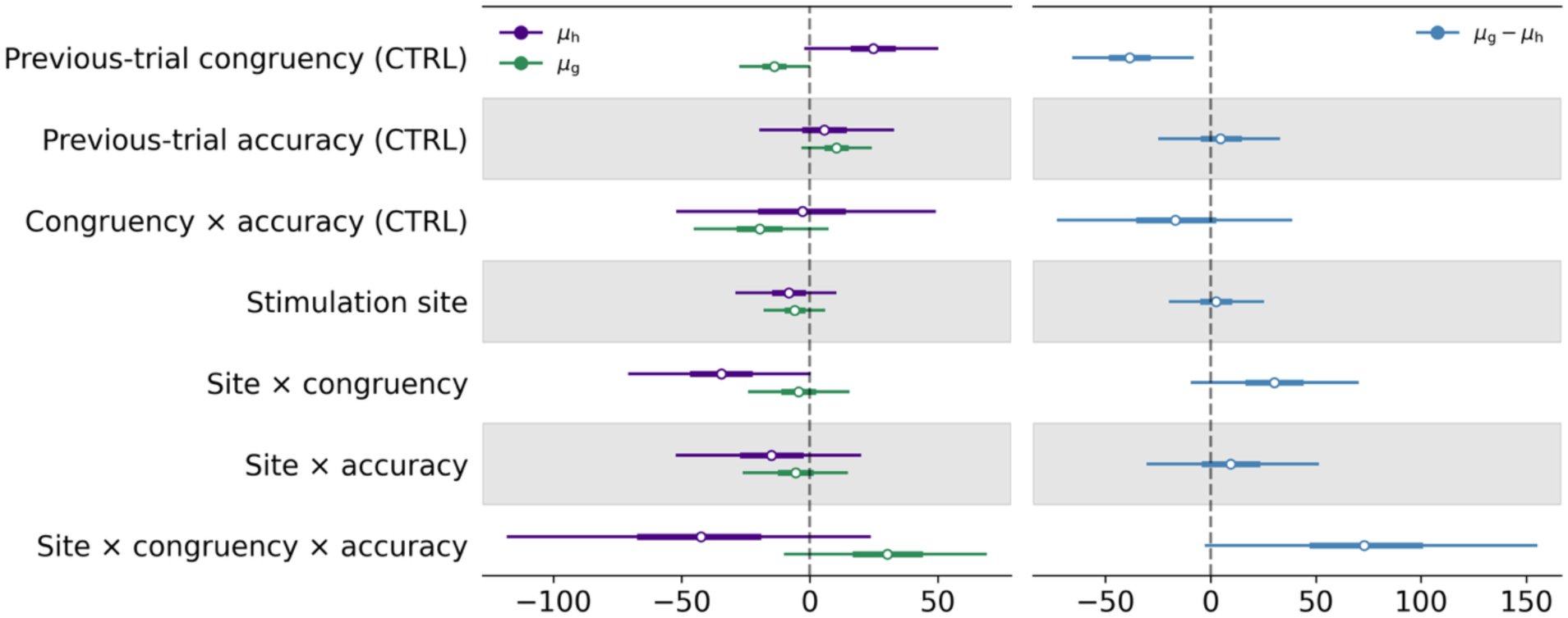
Effects of previous trial congruency, previous trial accuracy, and stimulation on habitual and goal-directed response preparation latency. Habitual responses (purple) were slower after an incongruent trial, and this slowing decreased following DLPFC stimulation. There was moderate evidence that this decrease in habitual response preparation latency after stimulation was even stronger following an error. Goal-directed responses (green) were prepared faster after an incongruent trial, and there was moderate evidence for a slowing of goal-directed response preparation following an error. There was moderate evidence this post-error slowing of goal-directed response preparation decreased following an incongruent trial. There was moderate evidence for a three-way interaction between stimulation site, previous trial congruency, and previous trial accuracy. We also computed the difference in goal-directed and habitual response preparation latency for each effect (blue). We observed strong evidence that this difference decreased following an incongruent trial, moderate evidence that DLPFC stimulation dampened the overall effect of previous trial congruency, and strong evidence DLPFC stimulation further dampened the effect of previous trial congruency after an error.

Next, we examined the effects of stimulation and trial history on goal-directed response preparation (i.e. processing of the task-relevant arrow direction; Figure 5, green). Following control stimulation, we found strong evidence that goal-directed responses were prepared faster after an incongruent trial (*b* = -13.7 ms, *95CI* = [-27.5, 0.251], *pd* = 0.972). We found moderate evidence that goal-directed responses were prepared more slowly after an error following control stimulation (*b* = 10.5 ms, *95CI* = [-3.10, 24.3], *pd* = 0.935). We found moderate evidence for an interaction between the CSE and the post-error effect (*b* = -19.5 ms, *95CI* = [-45.9, 6.95], *pd* = 0.927), suggesting that the speeding of goal-directed processing after an incongruent trial was further increased following an error. We found weak evidence for an effect of DLPFC stimulation on goal-directed response preparation (*b* = -5.79 ms, *95CI* = [-17.8, 6.30], *pd* = 0.828). We found weak evidence for an interaction between stimulation site and the CSE (*b* = -4.40 ms, *95CI* = [-24.3, 15.5], *pd* = 0.670). We also found weak evidence for an interaction between stimulation site and a post-error effect (*b* = -5.50 ms, *95CI* = [-26.2, 15.0], *pd* = 0.704). However, paralleling the effect on habitual response preparation, we found moderate evidence for a three-way interaction between stimulation site, the CSE, and the post-error effect (*b* = 30.4 ms, *95CI* = [-9.42, 70.0], *pd* = 0.934). This effect suggests that DLPFC stimulation reduced any additional speeding of goal-directed processing that an error induced after an incongruent trial relative to after a correct trial.

We also computed the difference in posterior samples between goal-directed response preparation latency and habitual response preparation latency *μ*_g_ − *μ*_ℎ_ for each combination of stimulation condition, previous trial congruency, and previous trial accuracy (Figure 5, blue). A smaller difference indicates stronger recruitment of control processes, as a smaller difference means less time when a habitual response can be expressed in the absence of a goal-directed response. Following control stimulation, we found strong evidence that this difference decreased after an incongruent trial relative to after a congruent trial (*b* = -38.5 ms, *95CI* = [-67.3, -9.34], *pd* = 0.994). We found weak evidence for a change in this difference following an error (*b* = 4.63 ms, *95CI* = [-25.0, 33.0], *pd* = 0.628), and weak evidence for an interaction between the post-error effect and CSE (*b* = -16.6 ms, *95CI* = [-72.6, 39.5], *pd* = 0.720). We found weak evidence for a main effect of DLPFC stimulation on the difference in response preparation latency (*b* = 2.44 ms, *95CI* = [-20.0, 25.2], *pd* = 0.586). However, we found moderate evidence for an interaction between stimulation site and the CSE (*b* = 30.2 ms, *95CI* = [-9.36, 70.3], *pd* = 0.928), suggesting DLPFC stimulation reduced the CSE relative to control stimulation. We observed weak evidence for an interaction between stimulation site and the post-error effect (*b* = 9.41 ms, *95CI* = [-31.9, 50.0], *pd* = 0.679). Reinforcing our findings for habitual and goal-directed response preparation, we found strong evidence for a three-way interaction between stimulation site, the CSE, and the post-error effect (*b* = 73.5 ms, *95CI* = [-5.69, 153], *pd* = 0.967).

To summarize, using control stimulation as a baseline, we found strong evidence for a slowing of habitual response preparation and speeding of goal-directed response preparation after an incongruent trial following stimulation of a control region. We found moderate evidence for a slowing of goal-directed processing after an error and moderate evidence that this slowing was decreased following an incongruent trial. We found little evidence for a main effect of stimulation on habitual and goal-directed response preparation, suggesting that DLPFC stimulation did not alter the overall congruency effect. However, we observed strong evidence that DLPFC stimulation decreased the slowing of habitual processing after an incongruent trial and moderate evidence that this slowing was further reduced after an error. We also observed moderate evidence that DLPFC stimulation reduced the additional speeding of goal-directed processing after an incongruent error trial relative to after a correct trial. Reinforcing these findings, we observed moderate evidence that DLPFC stimulation reduced the effect of previous trial congruency on the difference in habitual and goal-directed response preparation and strong evidence that this effect was further reduced following an error. These results highlight the causal role of DLPFC in using recent context to regulate control processes.

## Discussion

The recruitment of cognitive control processes involves a network of brain regions, including dorsolateral prefrontal cortex (DLPFC). DLPFC has been proposed to bias processing in sensory and motor regions to enable goal-directed behavior when there is an increased need for control (Carter & Van Veen, 2007; Miller & Cohen, 2001). However, there have been scant studies that have used methods that can determine the causal contribution of DLPFC to adaptation of cognitive control processes. Prior work has left it unclear if DLPFC is primarily involved in differentially biasing processing in task-irrelevant and task-relevant processing regions. Two indices of control adaptation are the congruency sequence effect (CSE) and the post-error effect. Both behavioral effects have been proposed to be the result of phasic control demands modulating LPFC activity (Carter & Van Veen, 2007; Kerns et al., 2004; Ridderinkhof et al., 2004). Using a novel combination of non-invasive brain stimulation, the forced-response paradigm, and computational modeling, we tested the causal role of DLPFC in control adaptation. This combination of methods allowed us to test the link between DLPFC activity and two common indices of control adaptation, while also revealing whether DLPFC activity differentially affects habitual or goal-directed response preparation. Disruption of DLPFC activity via transcranial magnetic stimulation (TMS) resulted in a reduction in the CSE, and the CSE was further reduced or even reversed following an error. These disruptions in control adaptation following DLPFC stimulation altered both habitual and goal-directed response preparation. These results provide causal evidence that right DLPFC supports top-down control by maintaining and using recent context (e.g., a previous trial) to alter task-irrelevant and task-relevant processing.

Theories of cognitive control allocation have proposed that both the CSE and the post-error effect can be explained by the detection of conflict on the current trial leading to an increase in the recruitment of control processes (Botvinick et al., 2001, 2004). Regions such as DLPFC are often implicated in implementing these adjustments of cognitive control. Prior functional MRI work has observed an increase in DLPFC activity following an incongruent trial (Egner & Hirsch, 2005; Kerns et al., 2004), and some work has observed an increase in LPFC activity following an error (Danielmeier et al., 2011; Kerns et al., 2004; King et al., 2010). However, Danielmeier et al. (2011) did not find a relationship in this post-error DLPFC activity to changes in sensory and motor cortex activity, making it unclear if DLPFC played a role in implementing control adjustments. Noninvasive brain stimulation studies have observed changes in the CSE following stimulation of DLPFC (Friehs et al., 2020; Muhle-Karbe et al., 2018). The present work aligns with prior work arguing for a causal role of DLPFC in implementing control adjustments. Prior TMS studies have primarily relied on one index of control adaptation, the CSE, when investigating the causal role of DLPFC in cognitive control processes. The use of only the CSE is understandable considering the low number of errors in free RT studies. The forced-response method allowed us to build on this prior work by using both previous trial congruency and accuracy in our analyses, providing causal evidence for a general contribution of DLPFC to behavioral indices of control adaptation.

The post-error adjustment disrupted by DLPFC stimulation in the present work is different than the post-error slowing of response times commonly observed in free RT work (Danielmeier & Ullsperger, 2011). We do observe moderate evidence for a slowing of goal-directed response preparation following an error. However, DLPFC stimulation affects the interaction between previous error and previous incongruent adjustments. Post-error slowing is often interpreted as heightened response caution following an error to prevent subsequent errors (Danielmeier & Ullsperger, 2011). That is, post-error slowing is often interpreted as a shift in the speed-accuracy tradeoff. The conflict monitoring theory proposes that response conflict following an error specifically leads to this slowing (Botvinick et al., 2001), and fMRI studies have found neural activity in cognitive control regions predictive of the magnitude of post-error slowing (Danielmeier et al., 2011; Kerns et al., 2004; King et al., 2010). The forced-response method controls for this adaptive post-error adjustment, as this method directly controls for differences in the speed-accuracy tradeoff between conditions (e.g., after an error vs after a correct trial). Adkins et al. (2024) found that errors were more likely after committing an error in a forced-response task, but this task did not induce response conflict. The post-error adjustment affected by DLPFC disruption is more aligned with a post-error reduction of interference, which has been observed in prior work (King et al., 2010; Ridderinkhof, 2002). This result also leads to the prediction that DLPFC activity should be higher after an error on an incongruent trial, and the magnitude of this activity may be predictive of the congruency effect.

Prior work has argued that the target of adaptive control processes are task-relevant processing regions, task-irrelevant processing regions, and a combination of both task-relevant and task-irrelevant processing regions. For example, when performing the arrow Simon task, increases in cognitive control could be achieved by suppressing location responses in motor cortex, suppressing location activity in visual cortex, or enhancing sensory activity related to the direction of the arrow. Neural evidence has been mixed, as studies have argued that the CSE is best described by changes in task-relevant processing (Egner & Hirsch, 2005) and task-irrelevant processing (Gheza et al., 2025). However, multiple computational modeling studies have argued that the CSE is best explained by changes in task-irrelevant, distractor processing (Gheza & Kool, 2025; Koob et al., 2023; Luo & Proctor, 2022; Ritz & Shenhav, 2023). Following an error, neuroimaging work has found evidence for adjustments in task-irrelevant and task-relevant sensory regions, along with suppression in motor regions (Danielmeier et al., 2011; King et al., 2010). It should again be noted that these adjustments in neural processing were primarily related to post-error slowing, although King et al. (2010) did find a relationship between post-error reductions in interference and processing in task-relevant sensory regions. A key advantage of the response preparation model applied to forced-response data is that this modeling approach can distinguish between changes in task-irrelevant processing and task-relevant processing (Sellers et al., 2026). Responses in free RT studies primarily come from time periods after control processes override the influence of task-irrelevant information. The forced-response method allows for direct observation of the time-dependent influences of task-irrelevant and task-relevant processing on behavior, and the response preparation model allows one to estimate the time course of habitual responses prepared to task-irrelevant information and goal-directed responses prepared to task-relevant information. The present study finds evidence for DLPFC implementing control adjustments by biasing task-irrelevant and task-relevant processing. While we cannot distinguish between biasing of sensory processing and motor responses, future fMRI studies should be able to determine if behavioral adjustments are related to suppression of irrelevant (i.e., habitual) motor responses or modulation of activity in sensory regions responsible for processing task-irrelevant and task-relevant information. We predict increases in DLPFC activity should be related to the degree of modulation in motor or sensory cortices and to the speed of habitual and goal-directed response preparation in the arrow Simon task.

One limitation of the present work is that the arrow Simon task does not control for stimulus and response repetitions from trial-to-trial. The CSE results must be interpreted with caution, as they could be driven by priming effects (Braem et al., 2019; Mayr et al., 2003). Stimulation of DLPFC still affected the CSE, and future work could determine if these effects are specific to priming, previous trial congruency, or a combination of priming and previous trial congruency. Furthermore, the forced-response method and response preparation model may be useful for decomposing how priming and previous trial conflict contribute to the CSE. For example, the effect of stimulus and response repetitions and the effect of previous trial congruency may selectively alter habitual and goal-directed processing.

To conclude, we provide causal evidence that right DLPFC uses recent context to implement adjustments in cognitive control. Disruptive stimulation of DLPFC attenuated effects of previous trial congruency and errors, and stimulation affected both task-irrelevant and task-relevant processing. This result provides novel support for theories that implicate DLPFC in implementing control adjustments on a trial-to-trial timescale.

